# Potential and limits of the evolutionary rescue of harvested food webs

**DOI:** 10.64898/2026.03.01.708823

**Authors:** Théo Villain, Jean-Christophe Poggiale, Anaëlle Peley, Nicolas Loeuille

## Abstract

Fishing deeply alters marine food webs structure and can drive the evolution of species traits, whether the species are directly targeted or not. Yet, studies rarely account for fisheries-induced evolution, and consequences are generally interpreted at the single-species level. Theory however predicts that eco-evolutionary dynamics within food webs can either promote biodiversity maintenance or accelerate its decline. In this study, we investigate how evolution affects the robustness of trophic networks under fishing pressure. Modifying evolution speed and the allocation of fishing effort across 458 structurally distinct allometric networks enables us to show that evolution most often enhances robustness. Network evolutionary response however becomes more variable (and possibly negative) as evolutionary rates increase and when fishing preferentially targets predators. By contrast, fishing strategies that concentrate effort on lower trophic levels, or distribute it more evenly, promote network persistence through evolutionary rescue while substantially reducing the risk of evolutionary collapse. Moreover, our results appear to be sensitive to the main forces governing ecological dynamics within the network such as competition or predation intensity. Finally, the consequences of network evolution differ across trophic levels. Evolution often drives the collapse of higher trophic levels while simultaneously promoting evolutionary rescue and enhancing diversity at lower levels through increased diversification, thereby generating a trade-off between vertical diversity (number of trophic levels) and total diversity. This highlights the importance of accounting for evolutionary dynamics and food web functioning in fisheries management, and suggests that reducing predator mortality may help prevent network evolutionary collapse.

## Introduction

Fisheries exert profound impacts on the persistence of marine populations (Jackson et al., 2001; Pauly et al., 1998). By extracting more than 8% of global aquatic primary production and up to 35% of that produced on continental shelves (Pauly and Christensen, 1995), fishing threatens the functioning of marine ecosystems and their resilience to additional anthropogenic pressures such as climate change (Sumaila and Tai, 2020). Although an increasing proportion of fishing grounds are now under management (FAO, 2024) fisheries management remains largely based on monospecific approaches (Kempf, 2010) and dependent on ecological approaches that once improved the status of managed stocks (Schaefer, 1954). However, neglecting species interactions and eco-evolutionary dynamics can compromise both system conservation (Skern-Mauritzen et al., 2016; Trijoulet et al., 2020; Wood et al., 2018) and fishery yields (Conover and Munch, 2002). Species interact through trophic and competitive relationships, and the removal of large numbers of individuals from a given trophic level may trigger cascading effects on lower levels (top-down effects, *e.g*., Baum and Worm, 2009; Frank et al., 2005; Pinnegar et al., 2000, Scheffer et al., 2005). Conversely, it may threaten the persistence of higher trophic levels through resource depletion (bottom-up effects, *e.g*., Cury et al., 2000, 2011; Smith et al., 2011). Likewise, fisheries-induced evolution (FIE; *e.g*., Heino et al., 2015) can proceed rapidly (Olsen et al., 2004) not only because fishing inherently imposes substantial mortality, but also because it acts as a direct selective force on traits of commercial interest (Allendorf et al., 2008; Darimont et al., 2009). Such evolutionary responses typically involve body-size reduction and changes in maturation/reproduction schedules (Olsen et al., 2004; Uusi-Heikkilä et al., 2015), ultimately undermining the regenerative capacity of stocks (Barneche et al., 2018; Enberg et al., 2009). It is therefore essential to integrate both ecosystem-level processes (Pikitch et al., 2004) and trait evolution (Kuparinen and Merilä, 2007; Stockwell et al., 2003) when evaluating the effects of fishing.

The consequences of fishing on food webs and on species evolution represent two relatively distinct branches of the literature, although some studies have attempted to bridge these aspects (*e.g*., Kuparinen et al., 2016; Nonaka and Kuparinen, 2023). In the face of disturbance, evolution can theoretically restore growth rates and enable populations to persist, a process known as *evolutionary rescue* (ER, Gomulkiewicz and Holt, 1995). This effect is supported by both empirical observations (Carlson et al., 2014; Vander Wal et al., 2013) and theoretical models (*e.g*., Osmond and De Mazancourt, 2013), raising hope for the maintenance of biodiversity under long-term pressures such as fishing (Carlson et al., 2014). However, two major limitations constrain evolutionary predictions at the network level:

i. evolution does not always result in the rescue even when considering a given species populations (evolutionary deterioration & suicide, Chapter 11, Ferriere et al., 2004)
ii. even when evolution exerts similar effects on two distinct species, their interactions may lead to unexpected outcomes (Loeuille, 2019)

In contrast to ER (*i*), some models suggest that gradual reductions in population density may be driven by evolution (*evolutionary deterioration*; Ferriere et al., 2004; Matsuda and Abrams, 1994). This mechanism has likely contributed to the collapse of cod stocks off Canada (Olsen et al., 2004; Swain, 2011). Evolution can also lead directly to population collapse (*evolutionary collapse*; Ferriere et al., 2004), or even extinction in cases of bistability (*evolutionary suicide*; Parvinen, 2005). Secondly (*ii*), evolution can generate contrasting outcomes at the community level, notably due to competition (De Mazancourt et al., 2008) and trophic interactions (Osmond et al., 2017). In predator-prey systems, such effects have been shown to lead to indirect ER (Yamamichi and Miner, 2015), particularly because changes in the densities or phenotypes of one species can induce ecological or evolutionary responses in the interacting species (Palkovacs et al., 2011; Palkovacs and Post, 2009; Troost et al., 2008; Walsh et al., 2012). The potential reciprocity of such changes promotes the emergence of eco-evolutionary feedback loops (Hočevar and Kuparinen, 2021; Jusufovski and Kuparinen, 2020), with variable consequences for fisheries (Edeline and Loeuille, 2021; Villain et al., Chapter 2). At the scale of food chains, evolutionary change in a single species can further trigger trophic cascades (Shackell et al., 2010) or even evolutionary cascades (Luhring and DeLong, 2020; Walsh et al., 2012). Altogether, these mechanisms considerably complicate our ability to predict the implications of evolutionary rescue, and more broadly, the effects of evolution on the persistence of species within trophic networks (Loeuille, 2019; Yacine et al., 2021).

In light of these complexities, the literature on network stability can provide additional insights into the eco-evolutionary responses of networks to fishing. Network robustness is defined as the capacity to maintain structure and functioning in the face of disturbance (Dunne et al., 2002). A first consistent finding from these studies is that while network size alone does not predict robustness to extinctions (Dunne et al., 2002; Keyes et al., 2024; McCann, 2000), robustness tends to increase with connectance (Dunne et al., 2002; Keyes et al., 2024). When connectance is high, species extinctions can be buffered by the redundancy of trophic contributions (Dunne et al., 2002; Sanders et al., 2018), thereby enhancing resistance to disturbance. Although these studies are not explicitly evolutionary, they suggest that under fishing pressure, if evolution promotes the loss of trophic interactions (due to trait mismatches), networks would become more fragile. Conversely, if evolution fosters the establishment or strengthening of trophic links, it could enhance network stability in the face of exploitation. A second consistent pattern emerges: the extinction of highly connected species within networks disproportionately promotes collapse compared to random extinctions (*rivet-like thresholds*; Dunne et al., 2004, 2002). In these networks, the highest number of connections occurs between intermediate trophic levels and top predators (Cohen and Briand, 1984; Dunne et al., 2004), which are also the species most affected by fishing (Essington et al., 2006; Pauly et al., 1998; Stevens, 2000). Consequently, fishing strategies under which evolution favors the persistence of predators may enhance network stability relative to alternative strategies. These findings echo calls for distributing fishing pressure across the network to maintain upper trophic levels, as advocated in the concept of balanced harvesting (Garcia et al., 2012; Zhou et al., 2010).

In this study, we aim to understand how evolution modulates the robustness of trophic networks under exploitation. Specifically, we analyze (*i*) how the rate of evolution influences network robustness in the face of fishing, (*ii*) whether the distribution of fishing effort within the network affects its eco-evolutionary persistence, and (*iii*) whether evolution can have contrasting effects across trophic levels (*evolutionary rescue* versus *evolutionary collapse*). To address these questions, we first generate networks through the evolution of fish body size, a key trait for characterizing life-history attributes and trophic interactions among individuals (Brose et al., 2006; Brown et al., 2004). To obtain the most general response possible, we explore a wide diversity of networks by varying the intensity of mechanisms such as competition, predation, and resource availability within the system (Allhoff et al., 2015; Brännström et al., 2011; Loeuille and Loreau, 2005). To assess the effect of evolution, we then simulate fishing on these networks and compare their robustness with and without evolutionary dynamics. When varying the rate of evolution (*i*), we expect it to enhance network persistence, notably by increasing phenotypic evolvability and the likelihood of evolutionary rescue. We also anticipate that evolution will have more favorable effects when fishing is distributed equally across different phenotypes (*ii*) because it reduces fishing pressure on particular phenotypes. Finally, we expect that prey evolution will buffer them against the effects of fishing, thereby allowing their predators to persist for longer (*iii*).

Our study investigates whether the concept of evolutionary rescue can be extended to the scale of trophic networks and promote the persistence of exploited marine ecosystems. While we show that evolution is most often positive when considering the total diversity of the network, it also typically accelerates the demise of upper trophic levels. Predator-focused fishing poses the greatest risk, as it strongly promotes evolutionary diversification of lower trophic levels, sometimes leading to evolutionary collapse of the system. Finally, network evolution typically accelerates predator extinctions (*evolutionary collapse*), even while promoting the persistence of other trophic levels (*evolutionary rescue*).

## Methods

We aim to understand how evolution affects the robustness of food webs to fishing. In this section, we first describe how networks are structured and emerge when body size governs interaction strength and metabolic rates (1). A wide diversity of networks is generated by combining parameters that modulate predation intensity, competition strength, and the amount of basal resource in the model (Brännström et al., 2011). For each parameter set, networks are generated under different mutation rates. We then select realistic networks by comparing their properties to empirical marine food webs (2). Each selected network is then subjected to exploitation (3), with fishing effort increasing linearly over time. Networks are harvested either with or without evolution, under different capture scenarios. Robustness is quantified (4) using the area under the curve of remaining species diversity through time (Pocock et al., 2012). For each case, comparing robustness with and without evolution (5) allows us to isolate the evolutionary effects on food web persistence under fishing pressure. Finally, we extend the analysis to the finer scale of trophic levels in order to assess whether responses differ across levels (6).

### 1) Eco-evolutionary model and emergence of size structured food webs

Our approach builds on the evolutionary emergence of size-structured food webs, based on previous works on this topic (*e.g*., Allhoff et al., 2015; Brännström et al., 2011; Loeuille and Loreau, 2005). In a nutshell, such eco-evolutionary models are based on phenotypic variations of body size due to mutation-selection processes, assuming that variations of body size lead to systematic changes in fecundity, mortality and ecological interactions (as detailed below). Under such hypotheses, starting from a single body size (*t* = 0, Figure 1B) consuming a basal resource (which does not evolve), the diversification process can lead to the emergence of complex food webs. At a given time, the system therefore includes a set of body sizes. Their dynamics build on a Lotka-Volterra model that incorporates competition, trophic interactions and fishing-induced mortality (eq. 2).

**Figure 1.**
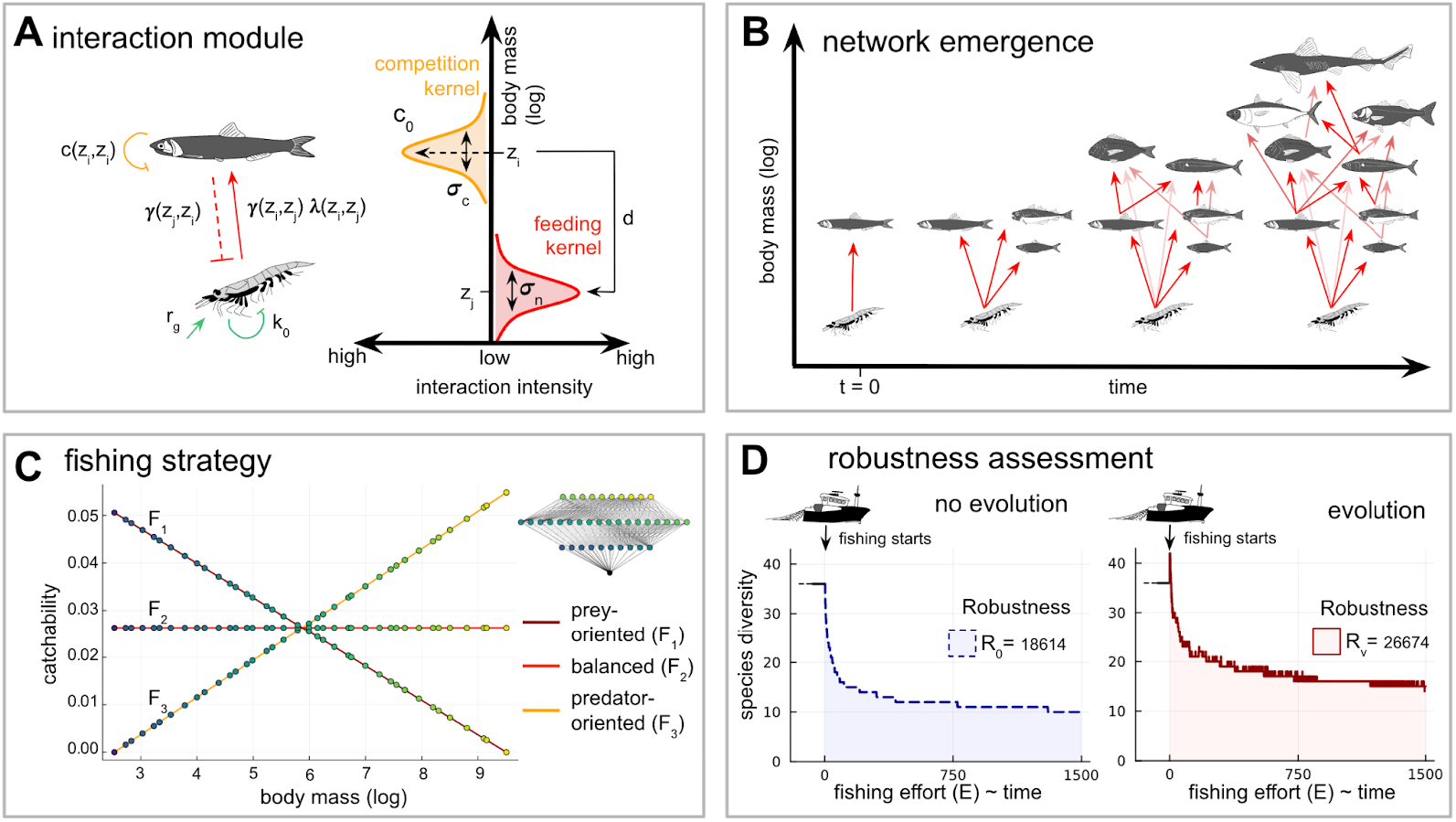
Generation of allometric food webs and robustness to fishing pressure. Interaction strengths depend on the body sizes of the different phenotypes (A). Each phenotype experiences competition with itself and with similarly sized individuals (niche of amplitude *c*_0_ and variance σ_*c*_), and preferentially consumes phenotypes that are smaller by a fixed size difference *d* (niche with variance σ_*n*_). B: The food web assembles through successive mutation–selection events, starting from an initial consumer–resource system (z(*t* = 0), where only the consumer evolves. C: After reaching an equilibrium, the resulting networks are subjected to three fishing strategies(*F*_1_, *F*_2_, *F*_3_), with or without evolution (D). D: Fishing effort increases linearly over time, and network robustness is assessed by integrating species diversity (as phenotypes that are close in trait space are grouped into a single species, see Methods 4.) over fishing effort, either with evolution (*R*_*v*_) or without (*R*_0_).

All generated food webs consist of a basal resource and consumers of diverse trophic levels:

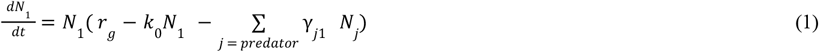

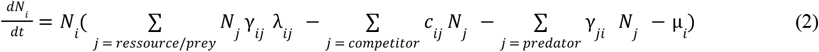

Where *N*_*i*_ represents the density of a population *i*. The strength of trophic and competitive interactions scales with allometric relationships between individual body sizes (Brännström et al., 2011). The size *z*_*i*_ of population *i* corresponds to the natural logarithm of adult body mass (Brose et al., 2008; Brown et al., 2004). The basal resource *N*_1_ sets the energetic constraints on which the food web is built and can be construed as a quantity of available inorganic nutrients. When no phenotypes are present, it follows logistic growth dynamics, reaching an equilibrium *r*_*g*_ / *k*_0_ (eq. 1). In line with many observations of feeding interactions (Brose et al., 2006; Li et al., 2023; Naisbit et al., 2011) we assume that predation is maximal when consumers are several times larger than their prey. Each phenotype *i* (excluding the resource) therefore consumes prey of size *z*_*j*_ with an interaction strength γ_*ij*_, which is higher when prey size is close to its preferred size (niche width σ_*n*_ ; see Supplementary 1 and Figure 1A). Energy assimilation occurs with an efficiency λ_*ij*_ (Figure 1A). It measures the fraction of biomass from prey *j* that can be used for the reproductive growth of predator *i* and assumes higher trophic transfer when prey size is larger (Supplementary 1). The model also assumes that competition is influenced by phenotypic similarity (in the tradition MacArthur and Levins, 1967), *e.g*., because similar body sizes use space and resources more similarly, therefore having more intense interference competition (*e.g*., Bowers and Brown, 1982). Each individual therefore interacts competitively with its own population and with neighboring phenotypes (*c*_*ij*_, niche width σ_*c*_; Supplementary 1 and Figure 1A). Finally, in line with the metabolic theory of ecology, we assume that changes in body size lead to variations in life-history traits, so that background mortality rate μ_*i*_ (Supplementary 1) that decreases with increasing body size (Brown et al., 2004).

Competition and predation niche width jointly control the structure of simulated food webs while primary productivity influences their overall size (Brännström et al., 2011, Allhoff et al., 2015, Loeuille and Loreau., 2005). To explore the relationship between evolution and robustness across a broad diversity of food web architectures, we generated three sets of networks (*Nw*_1_, *Nw*_2_ and *Nw*_3_) following the ecological dimensions explored by Brännström et al. (2011). Each network then arises from distinct combinations of ecological parameters. *Nw*_1_ networks vary in the baseline competition strength (*c*_0_) and competition niche width (σ_*c*_) (orange, Figure 1A), whereas *Nw*_2_ networks differ in both the predation niche width (σ_*n*_) and the competition niche width (σ_*c*_) (red and orange, Figure 1A). Finally, *Nw*_3_ networks vary with primary productivity, through changes in intrinsic growth rate (*r*_*g*_) and resource competition (*k*_0_) (green, Figure 1A).

In the first part of our analysis, we investigate the effect of evolutionary rate on network robustness to fishing by manipulating the mutation rate (μ) from μ = 10^−3^ to μ = 10^−1^. When evolution is fast, mutations occur frequently (every 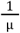 time step). Evolution is assumed to cease once a quasi-equilibrium is reached, defined as the point at which the weighted mean and variance of traits no longer exhibit significant changes, with absolute variations below 0.01.

### 2) Network selection

The vast parameter space we investigate (see above) leads to large variation in food web complexity. In particular, for some of the parameter combinations, species diversity is small (*e.g*., 2 or 3 species), and networks simple. Because we are here interested in the implications of ER in complex networks, we selected only networks whose structural properties closely match those of empirical marine food webs. For this selection, we extracted 169 trophic networks from the EcoBase database and calculated a set of 11 key properties: species richness, number of trophic links, connectance, proportion of basal, intermediate, and top predator species, maximum and mean trophic level, omnivory, generalism, and vulnerability. The same properties were computed for 1152 simulated networks and compared to the empirical ones using the Mahalanobis distance (MD; see Supplementary 2, Mahalanobis, 2018) to account for correlations among properties. Only the 458 networks whose structural features were closest to the empirical datasets were retained, using a threshold of MD = 40, which best differentiates MD values (Figure S2).

### 3) Network exploitation and fishing strategies

In the second part of the study, we analyze how the allocation of fishing pressure according to different strategies affects the evolutionary response of networks and changes their robustness. We assume a gradually increasing fishing intensity over time (eq. 3) to examine the transient response of the networks to harvesting. Incorporated into the ecological dynamic, phenotype *i* suffers from an additional fishing mortality rate − *q*_*i*_ *E*(*t*) which depends on both the fishing effort *E* and the selectivity *q*_*i*_ . The effort increases progressively until it reaches a maximum level (*E* = 1500) over 1. 5×10^5^ time steps (α = 10^−2^).

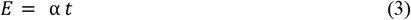

In ecosystem-based management, a key question is the understanding of how the distribution of fishing efforts affects the overall dynamics of the system. Some previous works for instance suggest that spreading evenly the fishing effort on species may be optimal. To tackle these questions, we here study a balanced harvesting scenario (designated as *F*_2_, see Figure 1C and Supplementary 3) as a baseline and contrast its results with two alternatives: a prey-oriented scenario (designated as *F*_1_, see Figure 1C and Supplementary 3) and a predator-oriented scenario (designated as *F*_3_, see Figure 1C and Supplementary 3). When exploitation begins, fishers distribute their effort across the different phenotypes, and the overall impact of fishing is identical for all networks and fishing strategies (eq. 4). This step thus defines the relationship between phenotype size and catchability in each network and remains fixed thereafter.

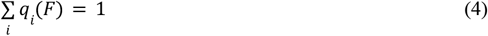

where *F* represents the fishing strategy.

### 4) Robustness assessment

To quantify the impact of fishing disturbance on networks, we use an approach based on their robustness to progressive species loss. Robustness is measured as the area under the curve (reviewed in Keyes et al., 2024) of remaining species diversity at each time step (as used in Pocock et al., 2012). In our study, the gradient of species loss is represented by fishing effort, and thus by time (eq. 3 & Figure 1D). Robustness (*R*) is therefore measured by integrating species diversity (*S*(*E*)) over the entire simulation (eq. 5).

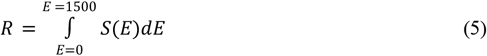

Since phenotypic diversity may not directly reflect species diversity, the trait space is discretized into intervals of width 2×σ_*z*_ (where σ_*z*_ corresponds to the mutation variance, see Table 1), and species diversity *S*(*t*) is computed as the number of occupied intervals.

**Table 1.**
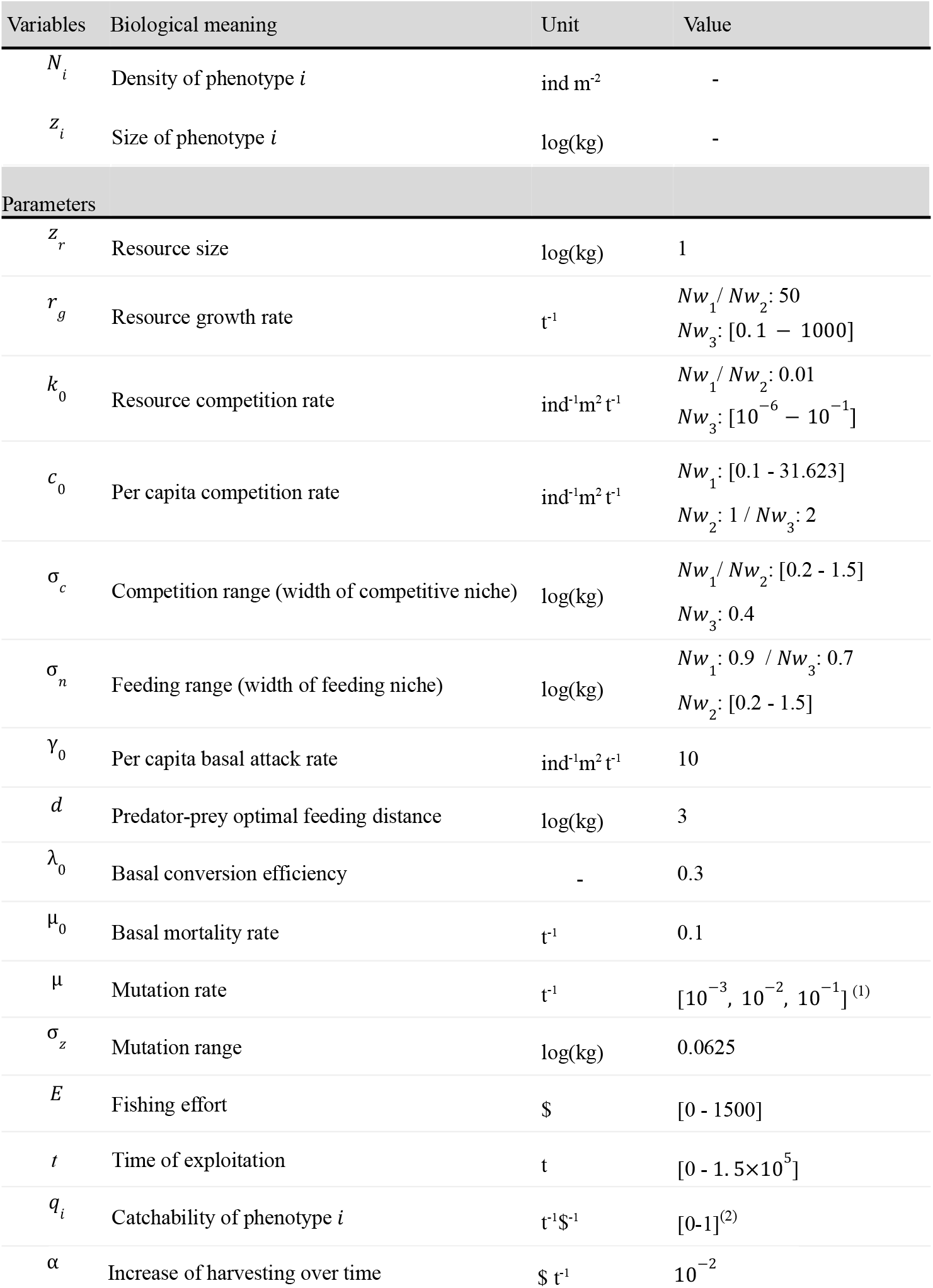
Variables and parameters used in the eco-evolutionary model. (1) changes when exploring the effects of the evolution rate. (2) changes when exploring the effects of the fishing strategy.

### 5) Effect of evolution

Each phenotype *i* has a size *z*_*i*_ that can evolve over time. To understand the effect of evolution on the robustness of a network under fishing pressure, we follow the same procedure as in Yacine et al. (2021). Networks are first subjected to a period without evolution for 1. 5×10^5^ time steps (grey in Figure 2A), so that counterselected mutants are excluded before the start of the fishing experiment. Each network is then exposed to fishing during 1. 5×10^5^ time steps either with or without evolution (dashed and solid lines respectively, as shown in Figure 2 and throughout the article). For each network, the robustness with evolution (*R*_*v*_) is then compared to the robustness without evolution (*R*_0_), allowing us to interpret the evolutionary impact on network robustness under fishing.

**Figure 2.**
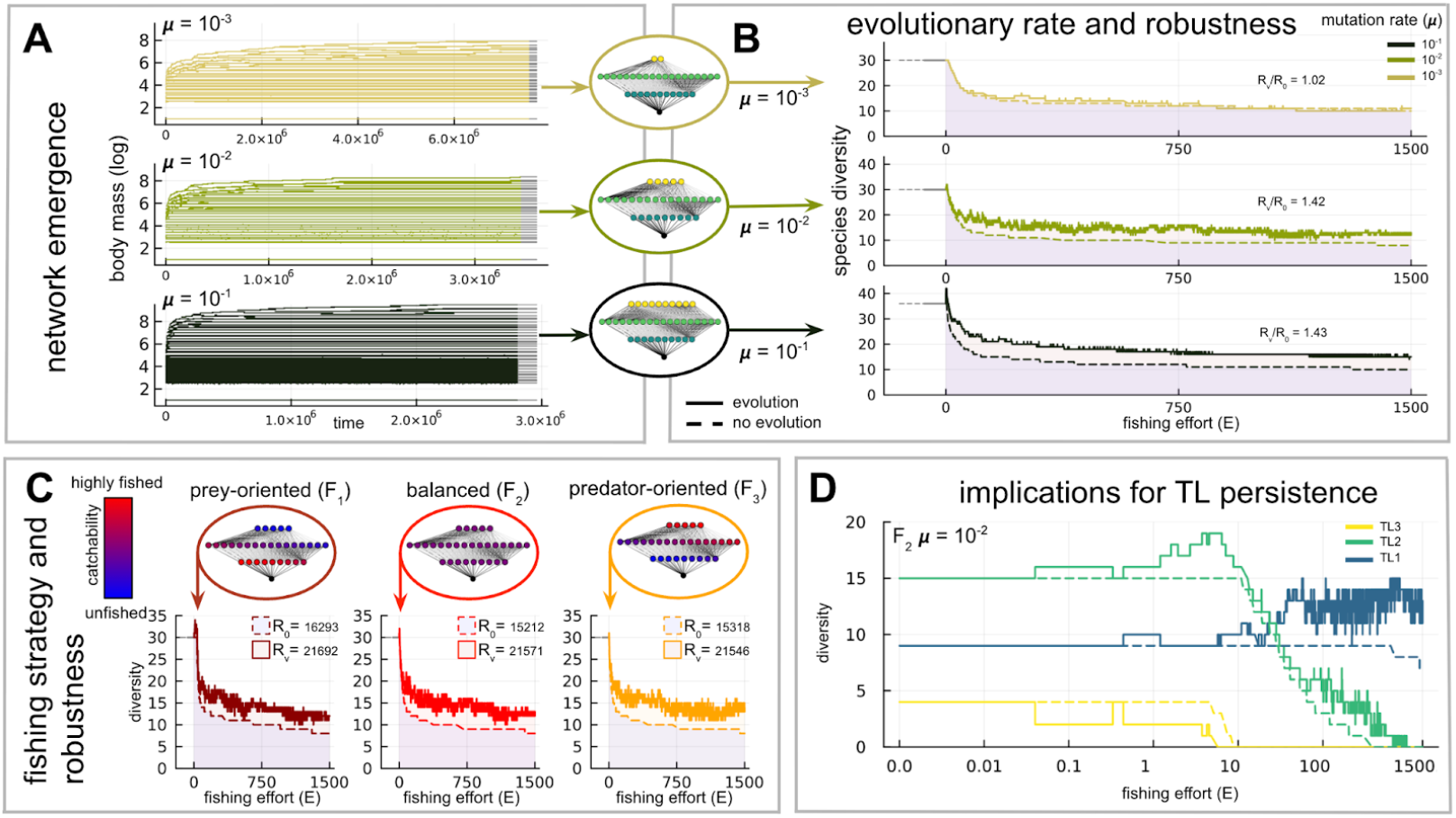
Effects of evolutionary speed and fishing strategy on network robustness. A-B-C-D: All networks share the same set of ecological parameters (networks from *Nw*_1_, *c*_0_ = 14. 678, σ_*c*_ = 0. 2). Each network is generated under different mutation rates (A). Once a quasi-equilibrium is reached, evolution is paused (grey) for 1. 5×10^5^ time steps. B-C-D: The networks are then subjected to harvesting, with fishing effort increasing linearly over time, and either with or without ongoing evolution (dashed or solid lines, respectively). *R*_*v*_ : robustness under exploitation with evolution. *R*_0_ : robustness without evolution. B: Effect of mutation rate on network robustness under fishing (balanced fishing condition *F*_2_). C: Effects of fishing strategy on the robustness of a network (mutation rate 10 ^−2^). D: trophic levels diversity under fishing pressure (balanced fishing condition *F*_2_, mutation rate 10 ^−2^).

### 6) Trophic level maintenance

Measuring robustness based on species diversity to quantify and interpret the evolutionary persistence of a network under fishing pressure may not always be appropriate. First, when networks evolve, their robustness to fishing can result from different mechanisms. Two limiting scenarios illustrate this: a network may exhibit a given robustness *R*_*v*_ because evolution promotes species persistence over time without leading to particular diversification (mutants systematically replace resident populations). The same robustness *R*_*v*_ may also arise because the network diversifies but collapses earlier. Second, when robustness is based on species diversity, it does not capture trophic reorganization. A network can display the same robustness *R*_*v*_ whether all top predators have gone extinct and lower trophic level species diversified, or conversely, if most species persisted without substantial diversification. Although robustness is equal in both cases, the conservation implications differ. To address these limitations inherent in our evolutionary approach, we measured the persistence or early disappearance of different trophic levels caused by evolution in the last part of the article (Figure 6). For each trophic level, we report the effort required to trigger its extinction with evolution, relative to that without evolution. When this ratio is greater than 1, network evolution promotes trophic level persistence (evolutionary rescue); when it is less than 1, it leads to earlier extinction of that trophic level (evolutionary deterioration or collapse).

## Results

We first detail the effect of evolution in response to fishing at the scale of a single characteristic food web. In the following sections, we then consider the full set of generated networks. The second section analyzes the effects of evolutionary rate on the evolutionary response, while the third section describes the consequences of the different fishing strategies. Finally, in the last section, we examine how these results at the network scale reflect the persistence of the different trophic levels.

### 1) In a given ecological system, both evolution speed and fishing strategies affect the robustness of food webs

At the scale of a food web, fishing consistently reduces species diversity, regardless of the scenario (with or without evolution), the speed of evolution, or the fishing strategy. Regarding the example of figure 2, when species evolve, robustness to fishing is systematically increased (*R*_*v*_ /*R*_0_ > 1), and this effect becomes more pronounced as the speed of evolution increases (Figure 2B). In the scenarios illustrated in Figure 2B, all species are fished with equal intensity (strategy *F*_2_),and fishing mortality is independent of body size (“balanced” harvesting). Evolution does not reduce fishing mortality directly, but can favor traits that improve reproductive success or reduce predation risk. As the speed of evolution increases, these effects are amplified: more mutants are generated at each time step, filling the trait space more densely. Under an increasing perturbation, when species have a greater evolvability they adapt more effectively and persist longer.

Preferential capture of certain species influences the robustness of the food web (Figure 2C). Fisheries primarily targeting prey species (strategy *F*_1_, dark red) improve robustness -with or without evolution-compared to other strategies. Indeed, without evolution the network has a robustness of *R*_0_ = 16293 using this effort repartition compared to *R*_0_ = 15212 and 15318 in balanced or predator-oriented strategies respectively, and the same results are obtained when the network evolves (*R*_*v*_ = 21692 compared to 21571 and 21546). However, the intrinsic effect of evolution is stronger when fishing is balanced or targets mainly predators (*R*_*v*_ /*R*_0_ increases by + 6. 5% and + 5. 6%, respectively). These counterintuitive differences arise from how the distribution of fishing effort affects the evolvability of different species. In prey-oriented fisheries, large-bodied (high trophic level) species are less frequently captured, resulting in higher densities. Because the fishing effort required for their extinction is higher, it explains the greater robustness of the network in the absence of evolution (dark red dashed line, *F*_1_). With evolution, higher densities of large species enhance their evolvability, allowing them to persist under greater fishing pressure. In contrast, smaller species are more heavily harvested, reducing their evolvability. Therefore, strategy *F*_1_ increases the relative evolutionary rate of predators compared to prey, promoting the maintenance of trophic transfer. Conversely, when top predators are more intensively exploited (strategies *F*_2_ and *F*_3_), fishing reduces predator evolvability and increases that of prey by altering their respective densities. As predators go extinct (e.g., TL3 in Figure 2D), prey species experience a sharp increase in evolvability due to the release from predation pressure. This boosts their densities and fosters diversification (e.g., around *E* = 7 Figure 2D), which explains why the evolutionary effect on robustness is stronger under strategies *F*_2_ and *F*_3_.

Finally, while evolution enhances the robustness of the whole network in this particular case, Figure 2D reveals that it can lead to both the persistence of certain trophic levels (evolutionary rescue) and the earlier loss of others (evolutionary deterioration). Indeed, comparing the solid and dashed lines for each trophic level shows that evolution promotes the extinction of TL3: diversity within this level drops to zero at a lower fishing effort than in the absence of evolution. This case of evolutionary deterioration contrasts with the evolutionary improvement observed for TL2, which persists under higher fishing pressure than without evolution.

Now that we have described possible dynamics and robustness constraints on an illustrative example, we assess whether the following qualitative outcomes are general. For simplicity, we only present the results from the harvesting of the realistic networks from the *Nw*_1_ parameters combinations, while commenting on the potential differences observed with the *Nw*_2_ and *Nw*_3_ networks. At this general scale, we ask whether the previous observations hold:

i. positive effects of evolution when considering the whole network diversity;
ii. enhancement of this positive effects in balanced and predator-oriented scenarios;
iii. evolutionary deterioration at upper trophic levels.

### 2) Trophic network diversity and the effect of evolutionary rate on robustness

Evolution has most often a positive effect when considering the overall diversity and increasingly enhances robustness as the evolutionary rate rises (Figure 3). At low mutation rates (μ = 10^−3^), however, evolution often has a negligible effect, and it can even reduce robustness (evolutionary deterioration in red, Figure 3B, *R*_*v*_ /*R*_0_ < 1) by causing early extinctions. These effects disappear at higher mutation rates, likely because early extinctions are offset by diversification of other lineages within the network. Moreover, since robustness depends on the number of species remaining over time, evolution can improve it through combined effects of species diversification or through evolutionary rescue that maintains evolving species under higher fishing efforts. Exploitation of the networks initially causes a surge in the diversity gap between scenarios with and without evolution (*S*_*v*_ (*E*)/*S*_0_ (*E*), Figure 3A), which is more pronounced at higher mutation rates (around *E* = 0 for all). As fishing effort increases, the diversity gap between the two scenarios continues to widen, and networks contain on average more species when evolution is allowed. Note that, regardless of mutation rate, evolution increases robustness continuously across the parameter space (Figure 3B-D). This effect appears to be clearly related to the intensity of competition within the network. As *c*_0_ increases, competition is stronger among populations of similar size. When σ_*c*_ increases, individuals can compete with populations that differ more in size. Increasing both parameters simultaneously further amplifies competition intensity. A certain *c*_0_ /σ_*c*_ ratio seems to promote network robustness. Conversely, when *c*_0_ is much lower than σ_*c*_, or the opposite, the evolutionary effects are less pronounced, and can even become negative at low mutation rates. Interestingly, high evolutionary effects occur on the most realistic networks (Figure S2). The consequences of evolutionary rates on networks evolutionary response to fishing are consistent with the *Nw*_2_ and *Nw*_3_ networks and extend even at lower evolutionary rates (Figures S4, S5, S6). Evolution favors networks persistence depending on the predation intensity (*Nw*_2_, Figure S5) and the total amount of basal energy (*Nw*_3_, Figure S6).

**Figure 3.**
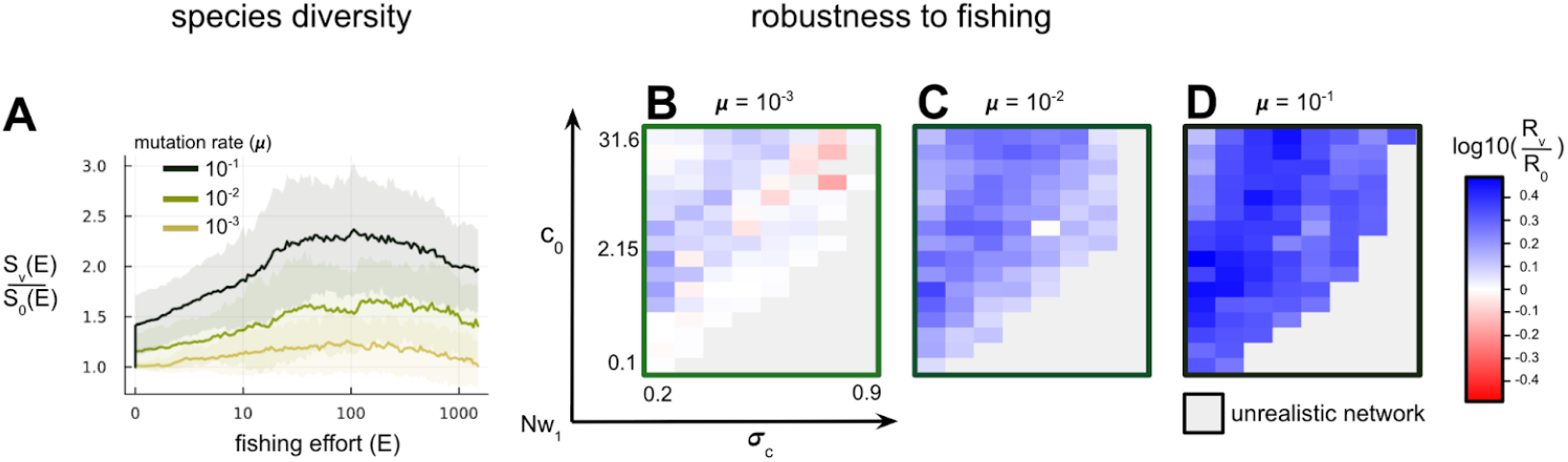
Evolution speed and the evolutionary maintenance of species diversity in response to fishing. For each mutation rate, all *Nw*_1_ networks are included, provided they meet the criteria for being realistic (see Methods). The networks are then subjected to fishing using strategy *F*_2_ . A: mean ratio between species diversity with evolution (*S*_*v*_) and without evolution (*S*_0_) as a function of fishing effort for different evolutionary rates. B-D: effect of the evolution on the robustness ratio (*R*_*v*_ /*R*_0_) under fishing pressure. For each set of ecological parameters, the effect of different evolutionary rates (from 10^−3^ to 10^−1^, from B to D) is tested. Blue: evolution increases network robustness. Red: evolution decreases network robustness. Grey: non-realistic network.

### 3) Trophic network diversity and the effect of fishing strategy on robustness

We now investigate whether the preferential harvesting of certain species alters the evolutionary responses of food webs. Overall, balanced or prey-oriented fishing leads to similar, positive effects of evolution, while results are more contrasted (including possible negative effects) when harvesting is predator-oriented. When large-bodied species are the main focus of fisheries (*F*_3_), evolution can both increase robustness and cause early network collapse (Figure 4). Because the relationship between fishing-induced mortality and fish size is positive (*F*_3_), smaller-sized mutants have a selective advantage over their resident population. Moreover, this fishing strategy greatly enhances the evolvability of small species because they experience both lower fishing pressure and reduced predation intensity compared to *F*_1_ and *F*_2_ . This can promote a rapid network reorganization and species evolution towards smaller sizes. In some cases, this transition may even lead to network collapse (in red Figure 4D). In other cases, diversification of small species helps maintain the underlying network. Although explaining the different outcomes remains difficult, it appears to be linked with competition again: when the niche breadth is relatively wide (intermediate values of σ_*c*_), small species favored by fishing exert stronger competitive pressure on larger species than when the competition niche is narrow (Figure 4B and Figure S7G for *Nw*_2_). Note also that networks that emerged under high predation intensity are more prone to collapse (Figure S7G), consistent with the effects of releasing strong top-down control when targeting predators. Conversely, balanced and prey-oriented strategies distribute fishing intensity more evenly across species thereby mitigating density imbalances. While the balanced (*F*_2_) scenario exerts no size-selective pressure, concentrating harvest on prey (*F*_1_) generates negative selection on body size and counteracts the benefits of species evolutionary shifts toward smaller sizes that would enhance productivity. The combination of these effects explains why these strategies limit the rapid reorganization of food webs and the risk of their collapse. Finally, the overall diversity maintained with evolution increases over time compared to non-evolving scenarios but do not really depend on the fishing strategies (Figures 4A, although contrasted for *Nw*_2_ and*Nw*_3_, see Figure S7).

**Figure 4.**
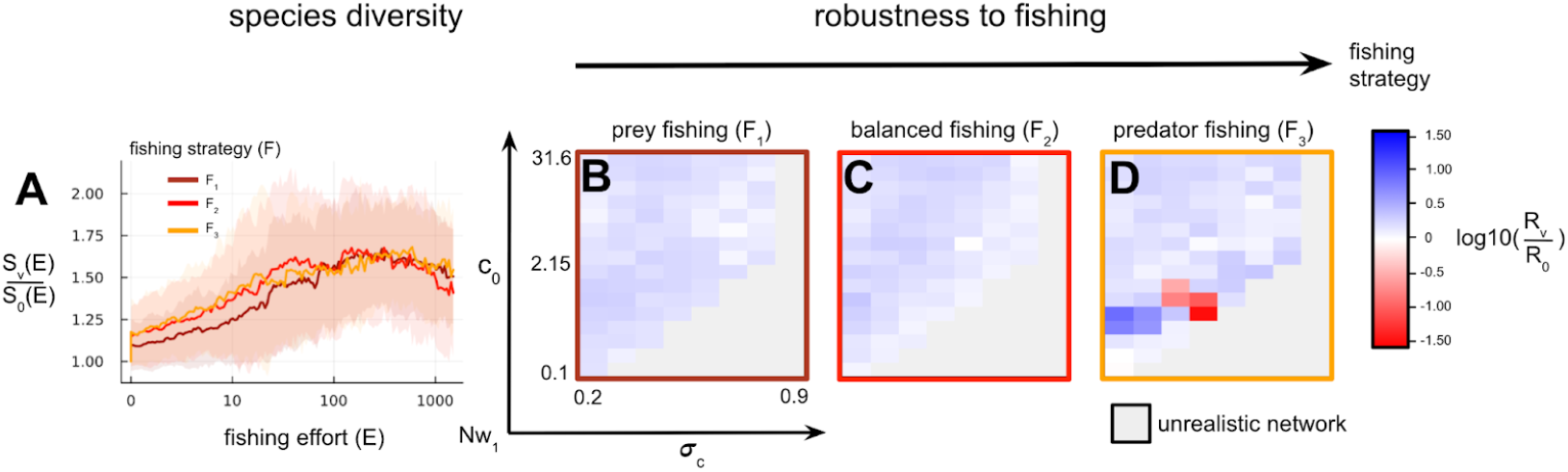
Effect of the fishing strategy on the evolutionary maintenance of species diversity in response to fishing. All *Nw*_1_ networks generated with a mutation rate of 10^−2^ are included, provided they are considered realistic (see Methods). A: mean ratio between species diversity with evolution (*S*_*v*_) and without evolution (*S*_0_) as a function of fishing effort for different strategies. B-C-D: effect of fishing strategy on robustness ratio (*R*_*v*_ /*R*_0_) in response to fishing. For each set of ecological parameters, the effects of different strategies (*F*_1_, *F*_2_, *F*_3_ from left to right) are tested. Blue: evolution increases network robustness. Red: evolution decreases network robustness. Gray: unrealistic network.

### 4) Evolutionary rate and predator-oriented fishing jointly increase the variability of evolution on robustness

After having separately examined the effects of fishing strategy and evolutionary rate across the different networks, we now analyze how these factors combine and influence the effect of evolution on fishing robustness (Figure 5). Regardless of the network, evolutionary rate generally promotes robustness across all fishing strategies (color by color in Figure 5), though note that negative outcomes are also possible (large error bars). When fishing predominantly targets prey, evolution less improves robustness compared to a balanced scenario (dark red vs. red in Figure 5). Predator-targeted fishing is characterized by greater variability in responses, and this variability increases with the evolutionary rate. Focusing captures on predators leads to more frequent and sometimes very early evolutionary collapses (*R*_*v*_ /*R*_0_ ≪1, even more pronounced in the *Nw*_2_ networks Figure S8B). Since networks that have emerged through intense predatory intensity (in *Nw*_2_ combinations) tend to face evolutionary collapse frequently under predator fishing, this strategy even leads to lower robustness compared to balanced or prey-oriented harvesting (Figure S8B). Our observations confirm previous suspicions: increasing fishing pressure on predators favors diversification of small-sized species. In some cases, this results in significantly greater network robustness, but in others, this diversification may precipitate network collapse.

**Figure 5.**
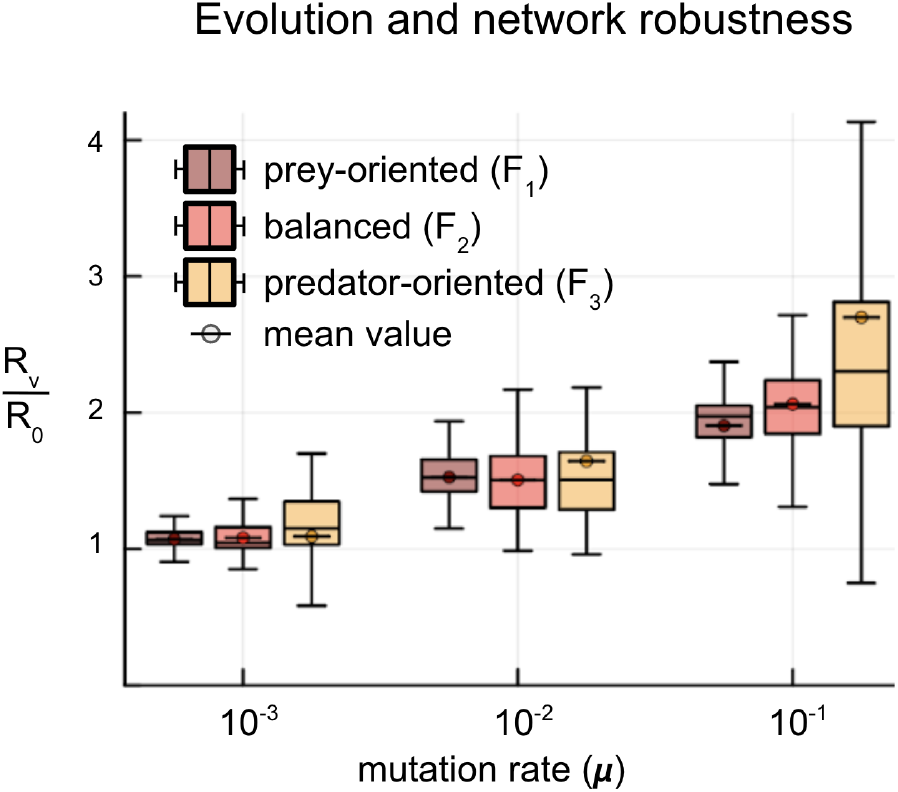
Combined effects of evolutionary rate and fishing strategy on the evolutionary persistence of trophic networks. Networks considered are *Nw*_1_ food webs. For each condition, boxplots show the distribution of values and means are indicated by circles.

### 5) Contrasting effects of evolution on the persistence of trophic levels

While the previous results may suggest that evolution is often positive, we stress that this is entirely due to the measure we take: the overall diversity of the network. Implications of evolution are even more contrasted when looking at changes in the network structure. We investigated this aspect by considering the role of evolution for each trophic level separately. Through evolution, a given trophic level can be maintained longer under fishing pressure (e.g., TL2 in Figure 2D) or, conversely, be driven more rapidly to extinction (e.g., TL3 in Figure 2D), even though the overall robustness increases (Figures 2B and 2C, μ = 10^−2^, *F*_2_). Figure 6 shows that this result is robust. Evolution often tends to accelerate the extinction of higher trophic levels (evolutionary deterioration) and to promote the persistence of lower trophic levels (evolutionary rescue). For instance under balanced harvesting, TL4 may collapse up to 100 times earlier compared to the non-evolving scenarios (Figure while TL2 persists longer. Comparing the effort required for extinction with (*E*_*v*_) and without (*E*_0_) evolution controls for the effect of diversification on overall robustness (since robustness depends on the number of remaining species at each time step) and provides additional insight into previous observations. Early diversity rebounds observed in Figure 3A&4A (around *E* = 0) may then be explained by the rapid disappearance of top trophic level species due to network evolution which further promotes diversification at lower trophic levels. Diversification would then explain the evolutionary effects on robustness patterns. However, it still remains difficult to determine whether the positive or detrimental effect of evolution on the persistence of one trophic level cascades to the lower trophic level. When evolutionary improvement is significant at a given level, it is generally associated with evolution accelerating the extinction of the upper trophic level (Figure S9B) although a causal link cannot be strongly established.

**Figure 6.**
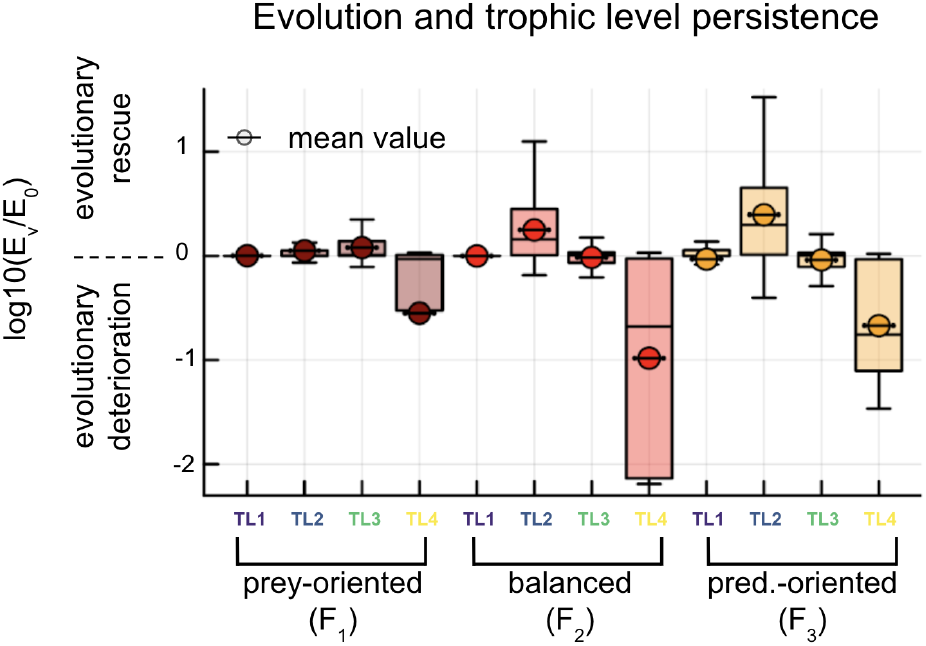
Evolutionary maintenance of different trophic levels as a function of fishing strategy. For each trophic level, the fishing effort leading to its extinction with evolution (*E*_*v*_) is compared to that without evolution (*E* _0_). Positive values indicate that evolution allows the trophic level to persist under higher fishing pressure (log10 scale). Negative values indicate that evolution causes the trophic level to collapse under lower fishing pressure. For each condition, mutation rates from μ = 10^−3^ to μ = 10^−1^ are taken into account. The basal resource TL0 is not considered since it is not harvested and does not evolve. Boxplots show the distribution of values, and means are indicated (circle).

Finally, the different fishing strategies also differentially affect the evolutionary maintenance of trophic levels (color by color, Figure 6). When fishing imposes low mortality on large-bodied (and thus high-trophic-level) species (*F*_1_), the evolutionary effect (positive or negative) on trophic level maintenance is weaker compared to other strategies. When predator mortality is higher (*F*_2_ and *F*_3_), evolutionary effects are both stronger and more variable, which is consistent with previous findings.

## Discussion

Our study first demonstrates that evolution generally enhances the robustness of trophic networks under fishing. However, when the rate of evolution or the release from predation pressure increases the evolvability of small-bodied fish, evolution can either promote early network decline or, conversely, substantially enhance its robustness. When fishing strategies maintain predator evolvability relative to their prey, evolution also improves network robustness, and its consequences are less risky, i.e., less likely to trigger network collapse. Finally, our study shows that higher trophic levels are generally subject to *evolutionary collapse* or *evolutionary murder*, whereas lower trophic levels are maintained longer through *evolutionary rescue*. Although it remains difficult to draw firm conclusions regarding the existence of evolutionary cascades, our results clearly indicate that the lower evolutionary potential of top predators relative to their prey constitutes an additional threat to their persistence and to network stability under fishing.

The general message of this study is that, when solely focusing on total diversity, evolution generally tends to enhance the robustness of trophic networks in response to disturbance, represented here by fishing pressure. This result is consistent with Yacine et al. (2021), who found that evolution in response to climate change promotes network persistence. In both our study and theirs, evolution can weaken networks under certain conditions. However, our results differ from those of Kuparinen et al. (2016) and Nonaka and Kuparinen (2023), who report that evolution in response to fishing destabilizes trophic networks. Several factors may explain these differences. First, the two studies quantify network stability using the temporal variability of fished fish biomass, whereas our study uses the area under the curve of species diversity. These metrics represent complementary but distinct facets of network stability (Domínguez-García et al., 2019). In their studies, evolution is not a stochastic response to fishing; age and size at maturity decline in a regular manner. Fishing acts as a constant pressure, whereas in our study it increases linearly over time. Finally, their models do not account for interspecific competition, whereas our results clearly depend on the intensity of competitive interactions.

To understand the implications of evolutionary rescue at the network level, we compared network robustness with and without evolution. This robustness-based approach allows us to integrate the evolutionary histories of all populations: when the evolutionary rescue events are frequent, evolution mitigates the negative effects of fishing and this is reflected in increased robustness. We find that, at the network scale, the rate of evolution generally enhances robustness. This result is not surprising, as one of the underlying assumptions of evolutionary rescue is that the frequency of mutation events promotes the emergence of better-adapted traits (Bell, 2017; Carlson et al., 2014). A second, complementary mechanism helps fully explain the effect on robustness: diversification. In response to environmental change and network reorganization, selection can be disruptive, maintaining new lineages (Rueffler et al., 2006), particularly under strong competition (Dieckmann and Doebeli, 1999). Diversification not only increases network robustness numerically but may also stabilize the network, allowing it to persist longer under perturbation (Yachi and Loreau, 1999). The effects of diversification have previously been predicted over the long term following periods of climate warming (Yacine et al., 2021), but our study extends this by showing that diversification can occur rapidly and accompany the disturbance. Surprisingly, some networks are penalized by slow evolution and strengthened when evolution is rapid (e.g., Figure 3). In isolated populations, slow evolution limits the likelihood of evolutionary rescue (Lynch and Lande, 1993; Carlson et al., 2014). However, it is difficult to conceive that evolutionary rate alone could produce such opposite outcomes (*evolutionary rescue* versus *collapse*). It is possible that these populations experience *evolutionary collapse* independently of the evolution of other species. At high mutation rates, network diversification may numerically compensate for robustness losses. Conversely, some populations may undergo *evolutionary murder*, particularly because, at these evolutionary rates, they evolve little or not at all relative to larger populations (Loeuille, 2019).

Our analyses on the effects of fishing strategies provide further insights into the pace of evolutionary dynamics. On average, evolution enhances robustness across all strategies, yet prey-focused or balanced fishing appears less risky than predator-focused strategies, which can yield variable outcomes. Our results first suggest that harvesting predators accelerates network evolution, in line with Luhring and DeLong (2020), who showed that predator removal triggers evolutionary changes in body sizes across other trophic levels. In our study, predator-oriented fisheries reduce predator densities compared to alternative strategies, consistent with empirical observations (Marshall et al., 2016) and theoretical predictions (Tromeur and Loeuille, 2017; Zhou and Smith, 2017). However, this outcome depends on the amount of prey harvested under alternative strategies and the relative strength of bottom-up and top-down controls (Villain et al. Chapter 1). Reduced predator densities lead to increased prey abundances due to relaxed predation pressure (Tromeur and Loeuille 2017; Villain et al., Chapter 1). Moreover, predator-targeted strategies exert weaker direct pressure on prey, further increasing their densities relative to other strategies. Finally, the relationship between fish size and fishing pressure is positive, which impacts population fitness, drives evolutionary shifts toward smaller body sizes, and consequently increases population densities (White et al., 2007). These three mechanisms converge to amplify the densities of lower trophic levels. As a result, network evolution is accelerated at lower levels (Bell 2017; Carlson et al., 2014), potentially fostering diversification, triggering evolutionary rescue (Hiltunen et al., 2014), or conversely leading to evolutionary collapse. Our study further suggests that variability in evolutionary outcomes is closely linked to competition. These findings resonate with other theoretical results: the balance between interspecific competition and density dependence (*c*_0_ in our study) influences the occurrence of evolutionary rescue (Osmond and De Mazancourt, 2013), particularly because competition slows down evolutionary responses (Johansson, 2008; Osmond and De Mazancourt, 2013). Diversification itself may also limit evolutionary rescue through competition (De Mazancourt et al., 2008), which may account for part of the observed variability in outcomes.

Finally, we show that while the consequences of evolution are globally positive for diversity, they are more contrasted across trophic levels. Network evolution tends to drive the earlier collapse of upper trophic levels, while maintaining intermediate and lower levels for longer periods. These patterns are observed under all fishing strategies, but their average magnitude is strongest under predator-focused fishing. This observation calls for a multi-criteria approach of network robustness as effects of evolution here often leads to trade-offs between total diversity and vertical diversity. Establishing a causal link between these outcomes (evolutionary collapse/murder at TLn leading to evolutionary rescue at TLn-1), or even a correlation (Figure S10), remains challenging. Nevertheless, our results clearly indicate that predators, owing to their lower evolutionary capacity (here mediated by density effects, as discussed above), are disadvantaged by the evolutionary response of their prey to fishing. This outcome is consistent with theoretical predictions (Loeuille, 2019), but contrasts observations of indirect evolutionary rescues in predator–prey models (Hiltunen and Becks, 2014; Northfield and Ives, 2013; Yamamichi and Miner, 2015). For instance, Yamamichi and Miner (2015) modeled prey and predator evolution under the assumption of a growth–defense trade-off in prey. When predators decline due to environmental disturbance, prey evolve reduced defense, which in turn rescues predators. In contrast, in our study, evolutionary changes in body size do not involve a growth–defense trade-off. Instead, prey evolve toward smaller body sizes primarily in response to fishing, and only secondarily to predator dynamics. Smaller sizes increase prey abundance (equivalent to higher growth rates), while simultaneously reducing predator feeding efficiency (equivalent to stronger defense). Because fishing is the main driver of evolution and predators evolve more slowly, smaller prey increasingly escape predation, which leads to premature predator extinctions (evolutionary murder). This explanation does not apply uniformly across all trophic levels, notably because basal resources do not evolve. At the bottom of the network, the downsizing of prey is constrained by the size of the basal resource (Villain et al., Chapter 2).

While the implications of our analysis are of general scope, they remain constrained by the assumptions of our model. We attempted to account for the diversity of possible responses by generating networks along axes known to diversify food web structures (Brännström et al., 2011; Allhoff et al., 2015; Loeuille and Loreau 2005). We then selected the networks that appeared most realistic by comparing them to empirical properties. This step likely underestimates the diversity of food webs, particularly because empirical nodes are typically constructed by aggregating distinct species (Christensen and Pauly, 1992; Dunne et al., 2004). Moreover, this type of eco-evolutionary model produces networks that resemble trophic chains, as interactions are constrained by a one-dimensional trait space (Fritsch et al., 2019). As a consequence, keystone species are absent and modularity is low, whereas both factors contribute to network robustness (Dunne et al., 2002; Smith et al., 2011). In addition, the use of the area under the species diversity curve is a useful metric to assess robustness (Keyes et al., 2024), but diversification within certain trophic levels complicates the interpretation of overall network responses. Finally, we assumed that fishing repartition does not adapt to the state of the network. Although global fishing pressure is increasing (Anticamara et al., 2011), stock depletion can lead to the closure of certain fisheries (e.g., Canadian cod) and to the redirection of harvesting efforts (Essington et al., 2006; Pauly et al., 1998).

This analysis contributes to understanding how evolution modifies the response of food webs to fishing and opens new avenues for research. First, it would be valuable to couple these analyses with implications for fisheries yields. When evolution enhances robustness by driving network phenotypes toward smaller body sizes, does this translate into improved yields? This outcome varies across studies, since some results suggest a decrease in yields (Barneche et al., 2018; Conover and Munch, 2002) while others indicate an increase (Eikeset et al., 2013). Second, our analysis highlights that evolution complicates the classical relationship to network robustness (e.g., area under the curve of species diversity, R50, etc.). While food webs can be made more robust through evolutionary rescue of their populations, our study shows that this may lead to contrasting functional consequences. A promising direction would be to move beyond cumulative metrics (directly or indirectly linked to the number of surviving populations) and instead prioritize transversal metrics more directly tied to network functioning (*e.g*., maximum or mean trophic level).

This study highlights that evolution can strengthen food webs in response to fishing, but that its consequences are detrimental for upper trophic levels. While such approaches can help understand and predict certain phenomena, they must above all be coupled with efforts to reduce the exploitation of marine food webs.

## Supplementary materials

### Supplementary 1: Life-history and interaction parameters

The life-history parameters and the strength of predatory interactions depend on relative prey and predator sizes (expressed as a log(biomass) *z*_*i*_). Mortality rate (eq. 1) is assumed to be decreasing with size (Brown et al. 2004):

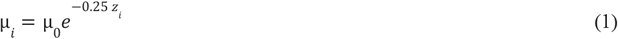

Attack rate is a function of predator-prey size ratio (eq. 2) with a maximal value when predators are *d* times larger than their prey (Brose et al. 2006) :

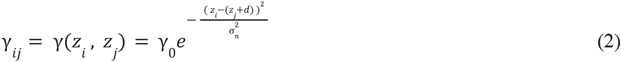

with *z*_*i*_ and *z*_*j*_ the respective predator and prey size. Note that predators can consume prey of increasingly diverse sizes as their trophic niche width σ_*n*_ increases. We assume a preferred predator-prey size difference *d* = 3 to be the same for all species. The conversion efficiency (eq. 3) is the efficiency of converting one unit of prey *j* into one unit of predator *i* and decreases with size difference:

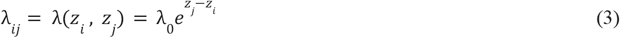

Interference competition between individuals is higher when sizes are close and decreases with size difference (eq. 4). The competition intensity between species from diverse sizes is higher when their competitive niche width σ_*c*_ increases:

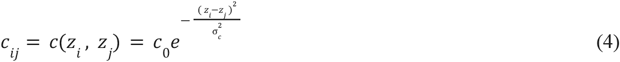

### Supplementary 2: Selection of realistic networks using Mahalanobis distance

To ensure the realism of the simulated networks, their properties were compared to those of 169 empirical food webs from the EcoBase database. Simulated networks whose properties most closely matched empirical ones were selected using the Mahalanobis distance (DM, Mahalanobis 1936), which accounts for the covariance structure among variables. The comparison was based on species richness, number of trophic interactions, connectance, the proportion of basal (resource-only consumers), intermediate, and top predator species, maximum and mean trophic level, omnivory, generalism, and network vulnerability. The properties of simulated networks were standardized based on the distribution of empirical networks (contained in *M*_*e*_), and aggregated into a matrix *M*_*S*_.

The covariance matrix of empirical network properties was then computed to reduce the influence of correlated variables in the Mahalanobis distance:

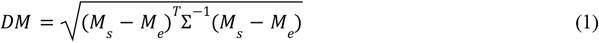

Each simulated network is then characterized by a DM that is lower when its properties are closer to those of the empirical food webs. Analysis of the resulting distances revealed a bimodal distribution, allowing the identification of a threshold (DM = 40) to distinguish the most realistic simulated networks from the others (Figure S2).

**Figure S2.**
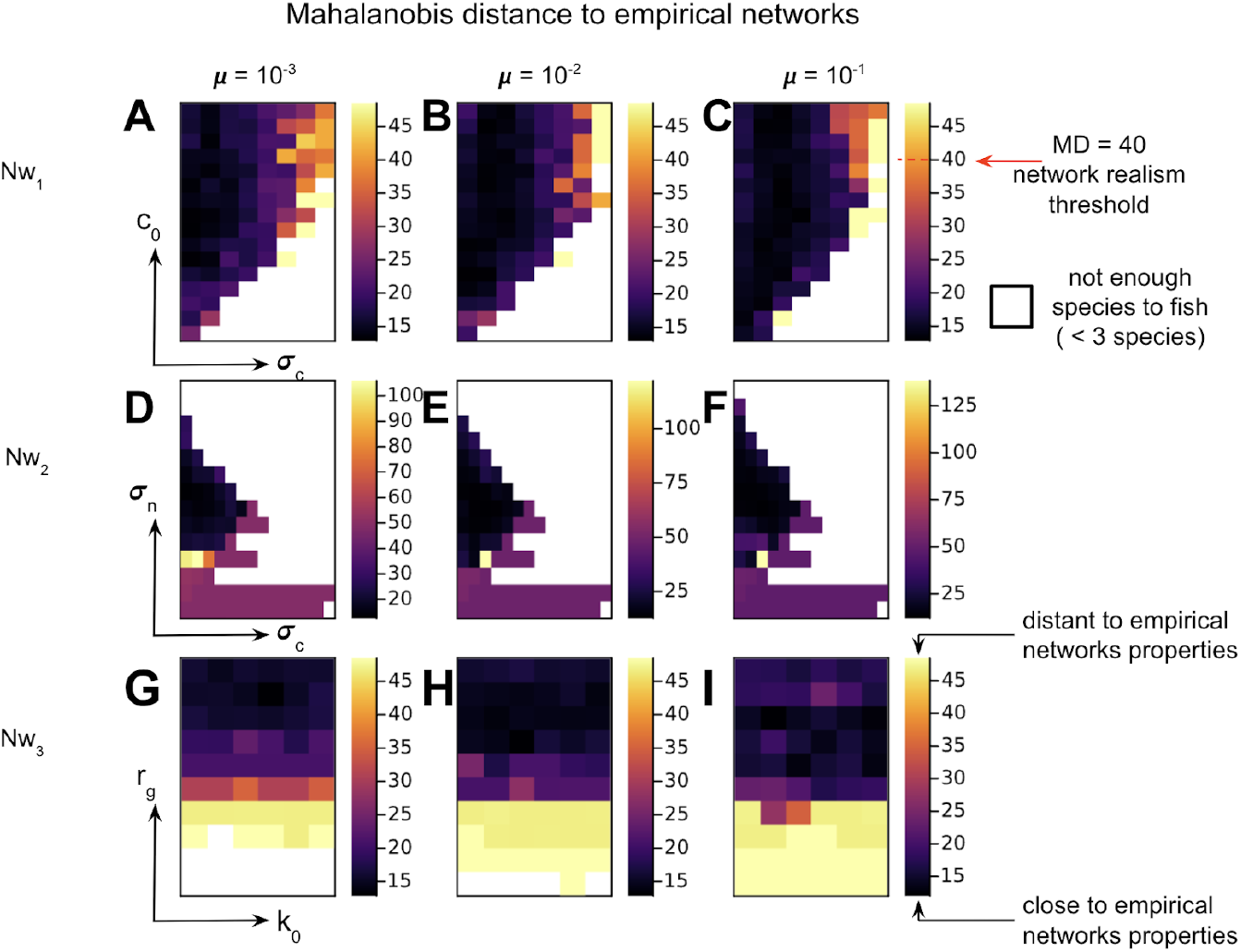
Mahalanobis distance between simulated and 169 empirical marine food webs from the EcoBase database. Eco-evolutionary networks were generated using the algorithm of Loeuille and Loreau (2005), followed by a 1. 5 ×10^5^ time-step period without evolution. Rows correspond to the three main types of simulated networks (from top to bottom: *Nw*_1_, *Nw*_2_, *Nw*_3_), and columns indicate the evolutionary rate (from left to right: μ = 10^−3^, 10^−2^, 10^−1^). Darker colors indicate greater similarity between simulated and empirical networks (i.e., lower Mahalanobis distance). White cells denote networks with fewer than three species (including the resource), which are insufficient to properly allocate fishing effort and are therefore excluded from further analyses.

### Supplementary 3: Fishing strategies

We aim to understand how preferential harvesting of certain phenotypes over others influences the eco-evolutionary dynamics of the system. We consider three fishing strategies, *F*_1_, *F*_2_, and *F*_3_ : *F*_1_ targets smaller phenotypes more intensely (prey-oriented fishing), *F*_2_ harvests all phenotypes equally (balanced fishing), and *F*_3_ targets larger phenotypes more intensely (predator-oriented fishing). In a network composed of *n* + 1 phenotypes (*n*≥2), the resource is assumed not to be harvested, and fishing effort is distributed among the *n* consumers. This distribution is based on the catchabilities *q*_*i*_ of the different phenotypes and allows for direct comparison between the different fishing strategies. The sum of catchabilities is then set to 1 regardless of the fishing strategy *F* (eq. 1).

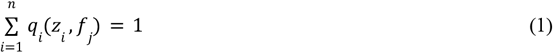

Where *f*_*j*_ is associated with fishing strategy *F*_*j*_ . It is proportional to the slope of the linear relationship (increasing, null, or decreasing, eq. 3) between catchability and phenotype size (eq. 2):

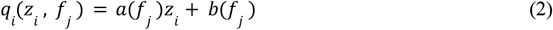

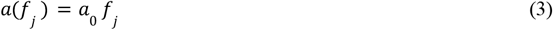

According to eq. 1, regardless of the phenotype considered:

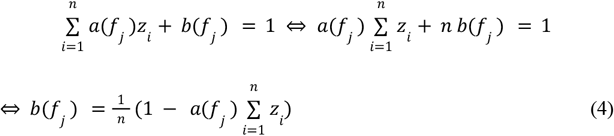

For clarity in the calculations, phenotypes are thereafter ordered by size (*z*_1_, *z*_2_,…, *z*_*n*_), with *q*_1_ representing the catchability of the smallest phenotype (of size *z*_1_) and *q*_*n*_ that of the largest phenotype (of size *z*_*n*_). Since catchability is modeled as an affine function of size, it is necessary to ensure that it remains non-negative. Accordingly, two limiting scenarios are defined based on the value of *f*_*j*_: when *f*_*j*_ > 0, the catchability of the smallest phenotype is set to zero; when *f*_*j*_ < 0, the catchability of the largest phenotype is set to zero. In the prey-oriented fishing strategy (*F*_1_), we consider the limiting case where the largest phenotype is not harvested (eq. 5):

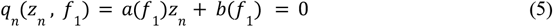

Then,

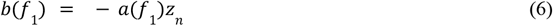

According to eq. 4 and eq.6,

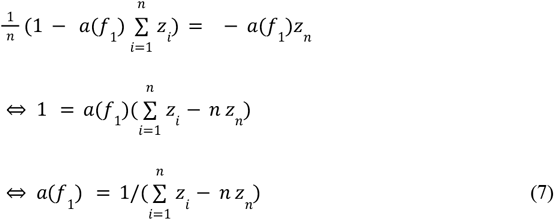

Similarly, for the predator-oriented fishing scenario (*F*_3_), we assume that the smallest phenotype is not harvested (limiting case, eq. 8), and we denote by *f*_3_ the corresponding slope:

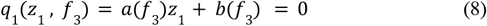

Thus,

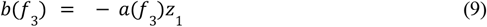

According to eq. 4 et eq.8,

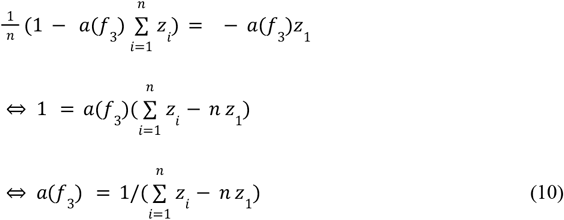

According to eq. 3,

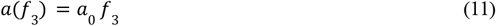

By setting *f*_3_ = 1, we obtain from eq. 11 that

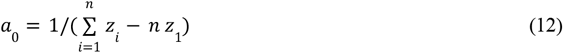

We note that *a*_0_ > 0 (eq. 13). Since there are phenotypes larger than the smallest one,

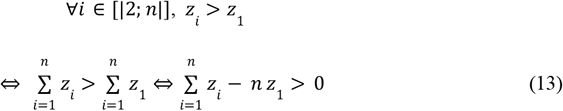

According to eq. 7, eq. 12 yields a value for the limiting slope associated with the fishing strategy *F*_1_ (eq. 14):

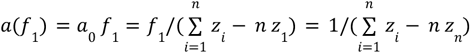

It follows that,

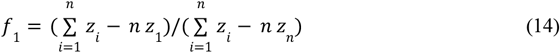

We also note that *f*_1_ < 0 (eq. 15). There exist phenotypes smaller than the largest phenotype:

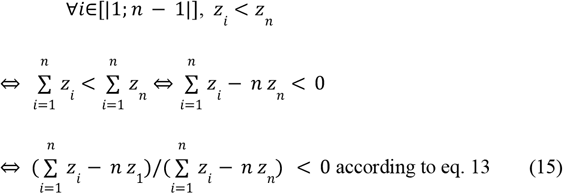

Therefore,

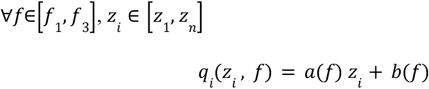

with

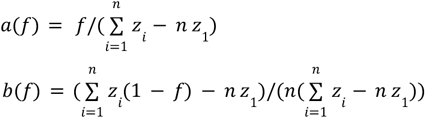

and

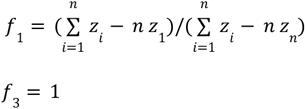

For simplicity, the fishing strategies are defined at *E* = 0 and remain unchanged thereafter. Fishers do not adapt their strategy based on potential extinctions of phenotypes or the emergence of mutants. Thus, when a mutant with size *z*_*m*_ appears during evolution, it will be fished only if its size lies between *z*_1_ and *z*_*n*_.

### Supplementary 4: Evolution speed (extended analysis) and the evolutionary maintenance of species diversity in response to fishing (*Nw*_1_ networks)

**Figure S4.**
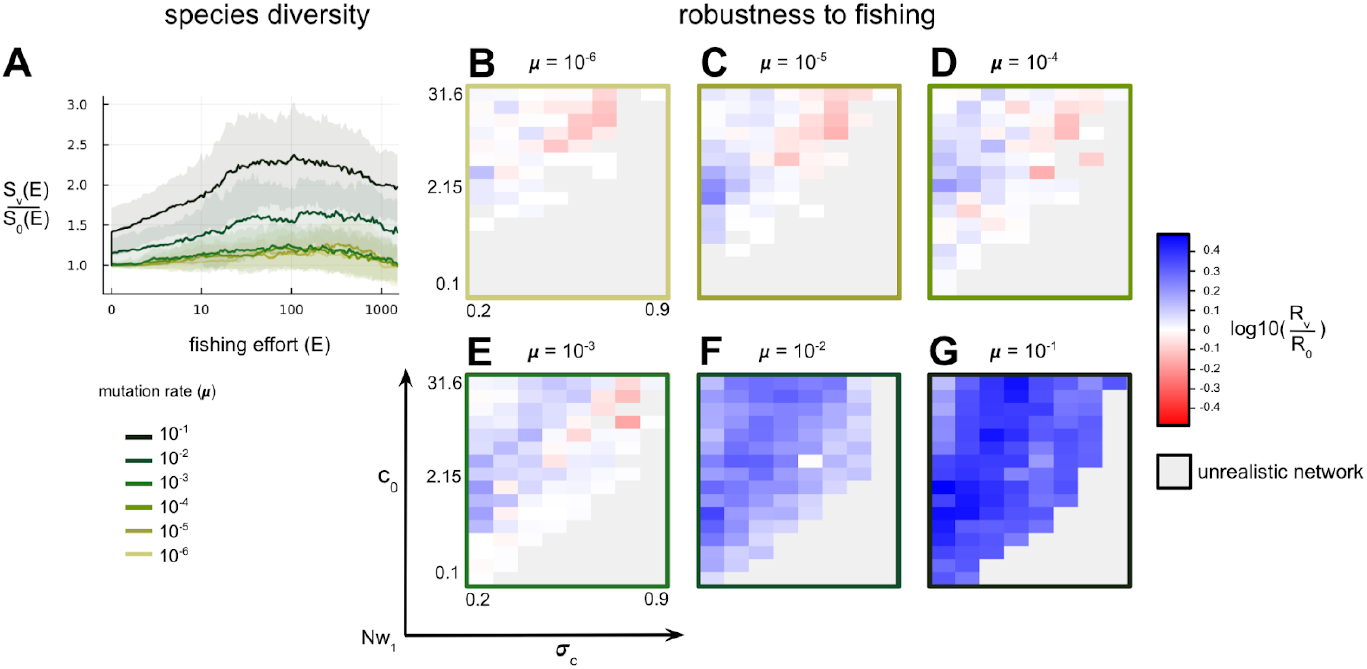
Evolution speed and the evolutionary maintenance of species diversity in response to fishing. For each mutation rate, all *Nw*_1_ networks are included, provided they meet the criteria for being realistic (see Methods). The networks are then subjected to fishing using strategy *F*_2_. A: mean ratio between species diversity with evolution (*S*_*v*_) and without evolution (*S*_0_) as a function of fishing effort for different evolutionary rates. B-G: effect of evolution on the robustness ratio (*R*_*v*_ /*R*_0_) under fishing pressure. For each set of ecological parameters, the effect of different evolutionary rates (from 10^−6^ to 10^−1^, from B to G) is tested. Blue: evolution increases network robustness. Red: evolution decreases network robustness. Grey: non-realistic network.

### Supplementary 5: Evolution speed and the evolutionary maintenance of species diversity in response to fishing (*Nw*_2_ networks)

**Figure S5.**
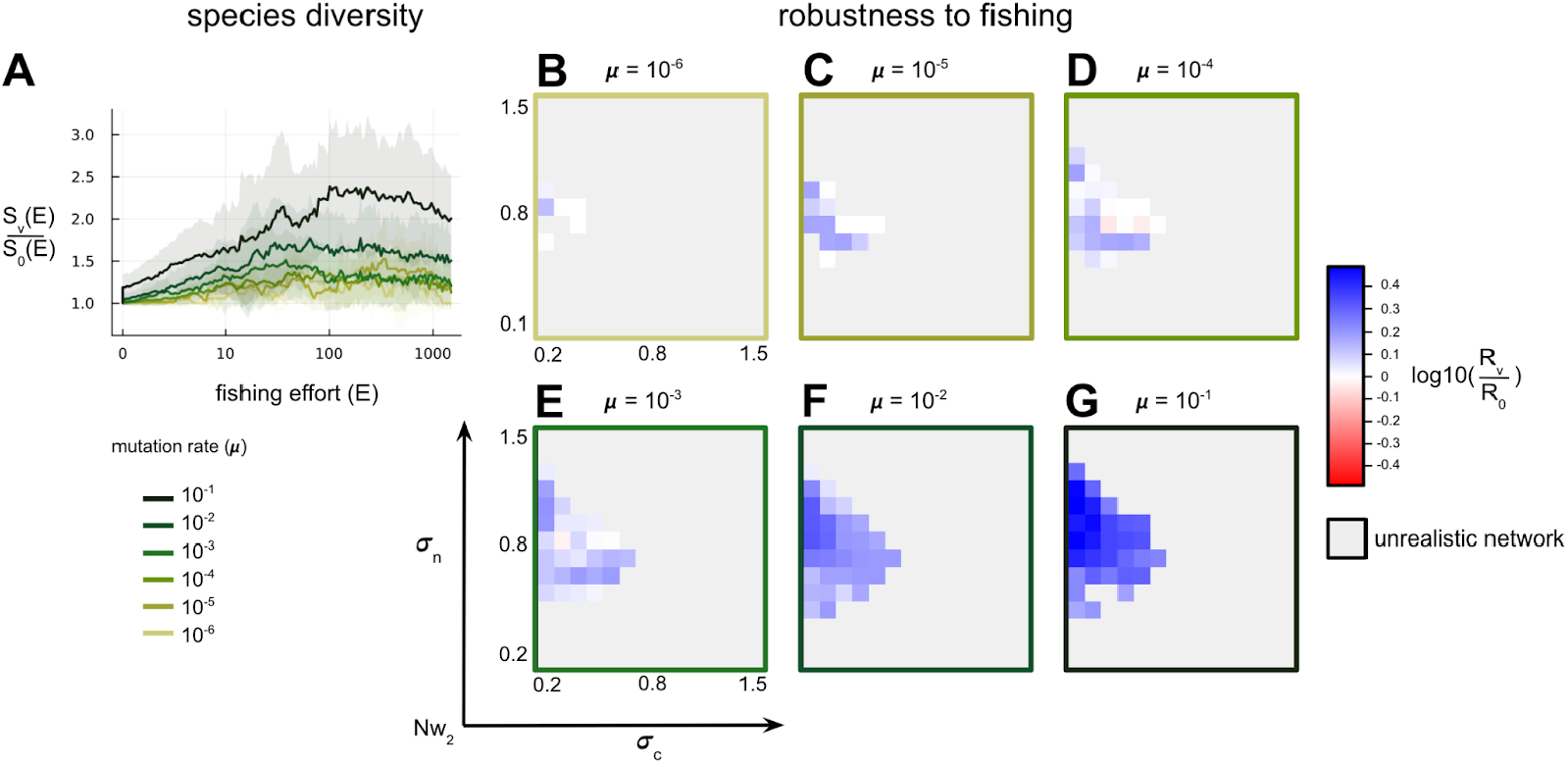
Evolution speed and the evolutionary maintenance of species diversity in response to fishing. For each mutation rate, all *Nw*_2_ networks are included, provided they meet the criteria for being realistic (see Methods). The networks are then subjected to fishing using strategy *F*_2_. A: mean ratio between species diversity with evolution (*S*_*v*_) and without evolution (*S*_0_) as a function of fishing effort for different evolutionary rates. B-G: effect of evolution on the robustness ratio (*R*_*v*_ /*R*_0_) under fishing pressure. For each set of ecological parameters, the effect of different evolutionary rates (from 10^−6^ to 10^−1^, from B to G) is tested. Blue: evolution increases network robustness. Red: evolution decreases network robustness. Grey: non-realistic network.

### Supplementary 6: Evolution speed (extended analysis) and the evolutionary maintenance of species diversity in response to fishing (*Nw*_3_ networks)

**Figure S6.**
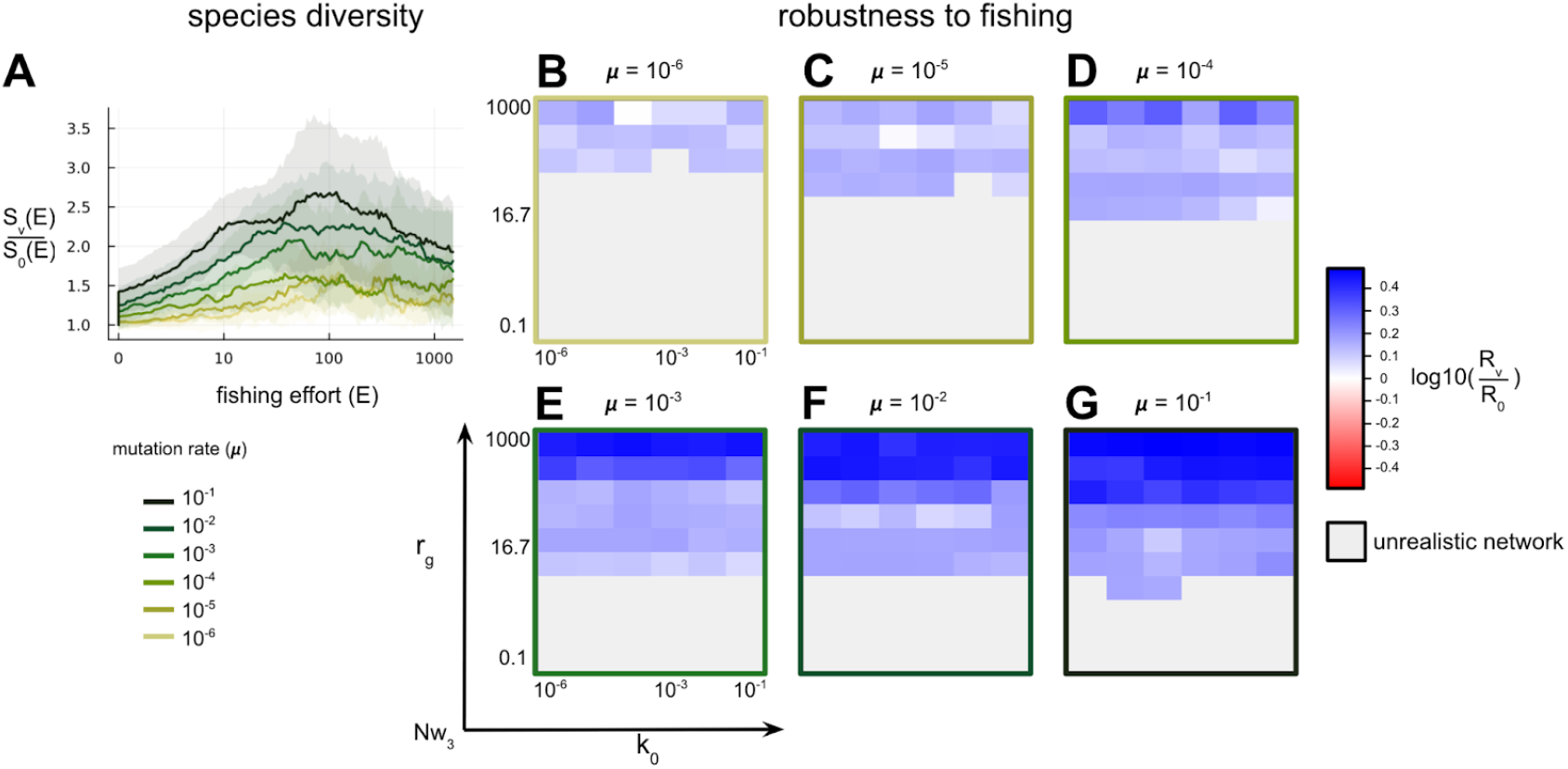
Evolution speed and the evolutionary maintenance of species diversity in response to fishing. For each mutation rate, all *Nw*_3_ networks are included, provided they meet the criteria for being realistic (see Methods). The networks are then subjected to fishing using strategy *F*_2_. A: mean ratio between species diversity with evolution (*S*_*v*_) and without evolution (*S*_0_) as a function of fishing effort for different evolutionary rates. B-G: effect of evolution on the robustness ratio (*R*_*v*_ /*R*_0_) under fishing pressure. For each set of ecological parameters, the effect of different evolutionary rates (from 10^−6^ to 10^−1^, from B to G) is tested. Blue: evolution increases network robustness. Red: evolution decreases network robustness. Grey: non-realistic network.

### Supplementary 7: Fishing strategy and the evolutionary maintenance of species diversity in response to fishing (all networks)

**Figure S7.**
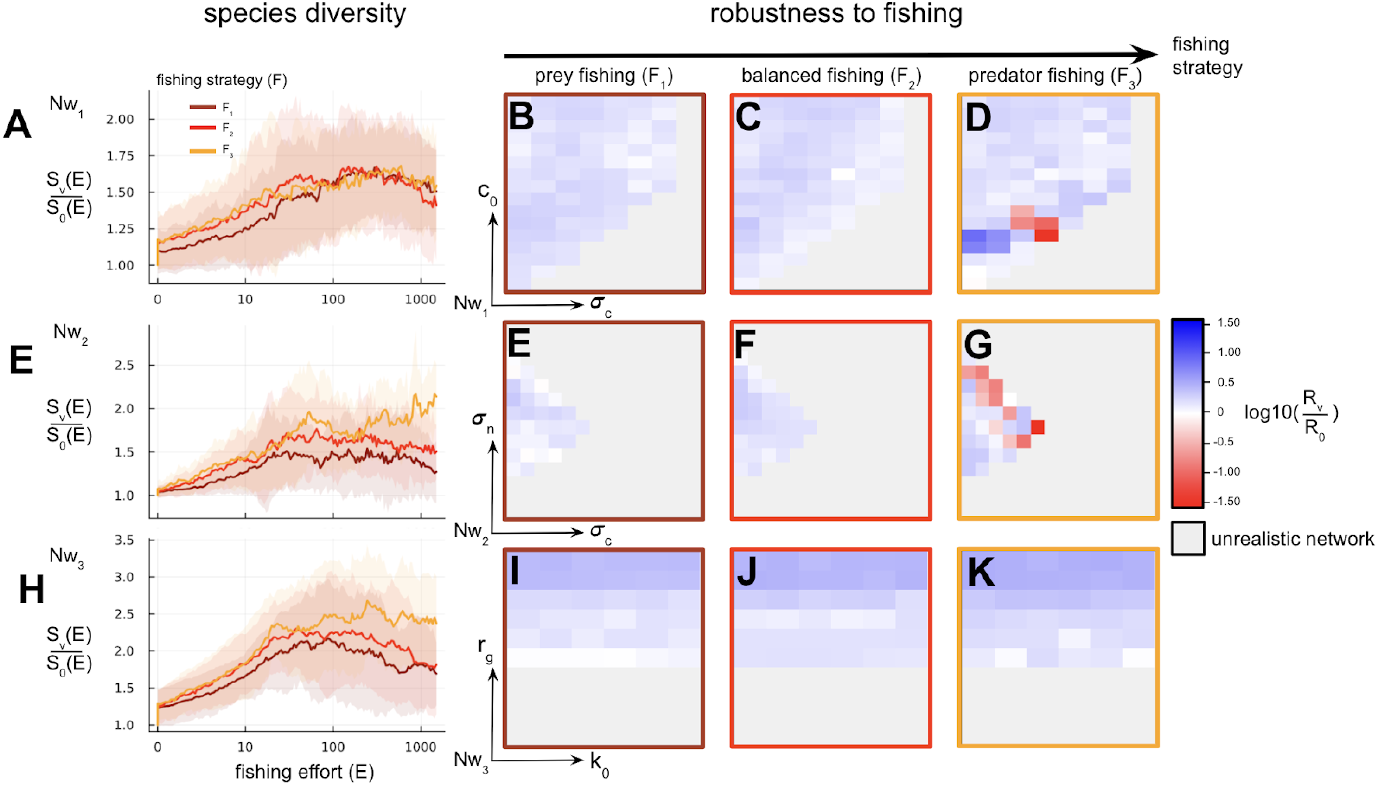
Effect of fishing strategy on the evolutionary maintenance of species diversity in response to fishing. All networks generated with a mutation rate of 10^−2^ are included (*Nw*_1_, *Nw*_2_, *Nw*_3_), provided they are considered realistic (see Methods). A-E-H: mean ratio between species diversity with evolution (*S*_*v*_) and without evolution (*S*_0_) as a function of fishing effort for different strategies. A-D: *Nw*_1_ networks. E-G: *Nw*_2_ networks. H-K: *Nw*_3_ networks. B-C-D/E-F-G/I-J-K: effect of fishing strategy on robustness ratio (*R*_*v*_ /*R*_0_) in response to fishing. For each set of ecological parameters, the effects of different strategies (*F*_1_, *F*_2_, *F*_3_ from left to right) are tested. Blue: evolution increases network robustness. Red: evolution decreases network robustness. Gray: unrealistic network.

### Supplementary 8: Fishing strategy and the evolutionary maintenance of species diversity in response to fishing (all networks)

**Figure S8.**
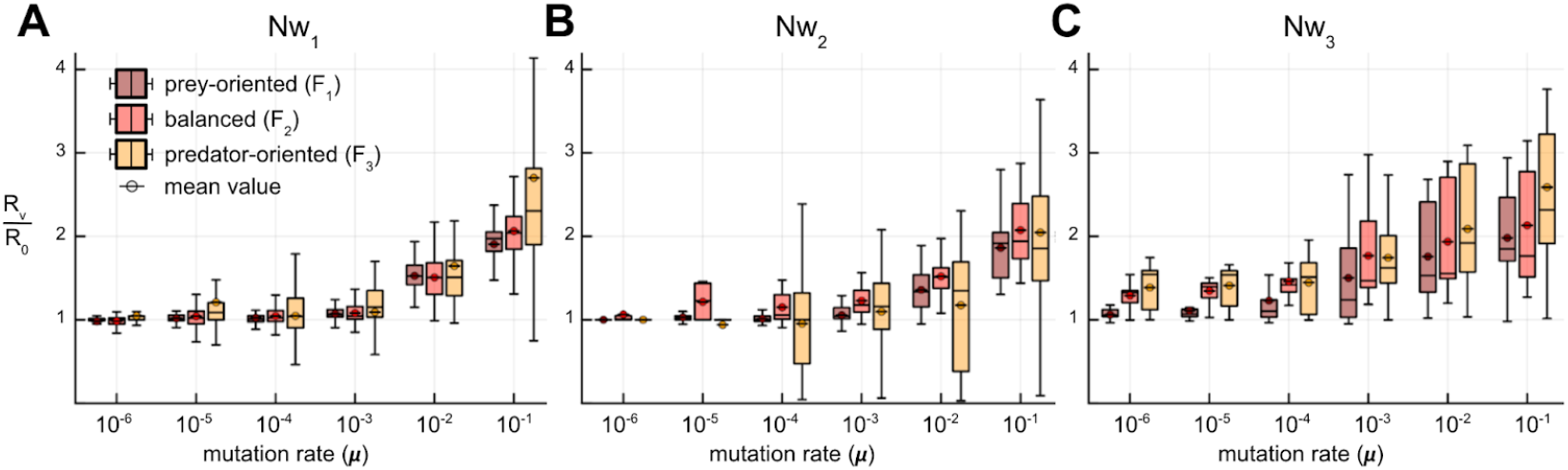
Combined effects of evolutionary rate and fishing strategy on the evolutionary persistence of trophic networks . A: Networks *Nw*_1_. B: Networks *Nw*_2_. C: Networks *Nw*_3_. For each condition, boxplots show the distribution of values and means are indicated by circles.

### Supplementary 9: Fishing strategy and the evolutionary maintenance of species diversity in response to fishing (all networks)

**Figure S9.**
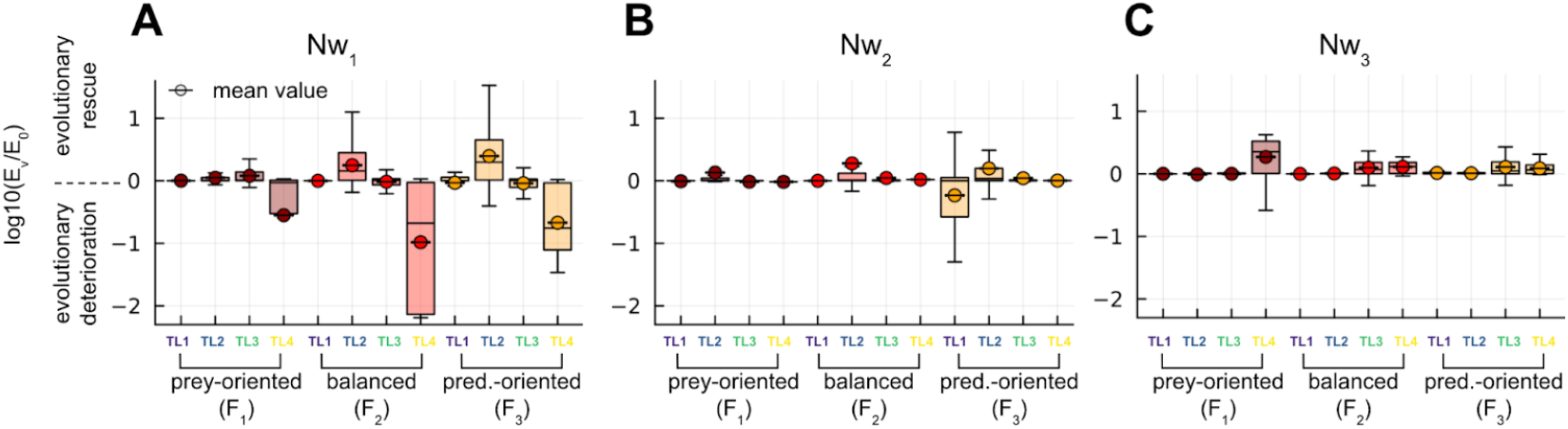
Evolutionary maintenance of different trophic levels as a function of fishing strategy. For each trophic level, the fishing effort leading to its extinction with evolution (*E*_*v*_) is compared to that without evolution (*E*_0_). Positive values indicate that evolution allows the trophic level to persist under higher fishing pressure (log10 scale). Negative values indicate that evolution causes the trophic level to collapse under lower fishing pressure. A: *Nw*_1_ networks. B: *Nw*_2_ networks. C: *Nw*_3_ networks. For each condition, mutation rates from μ = 10^−3^ to μ = 10^−1^ are taken into account. The basal resource TL0 is not considered since it is not harvested and does not evolve. Boxplots show the distribution of values, and means are indicated (circle).

### Supplementary 10: Evolutionary events and cross-trophic level relationships

**Figure S10.**
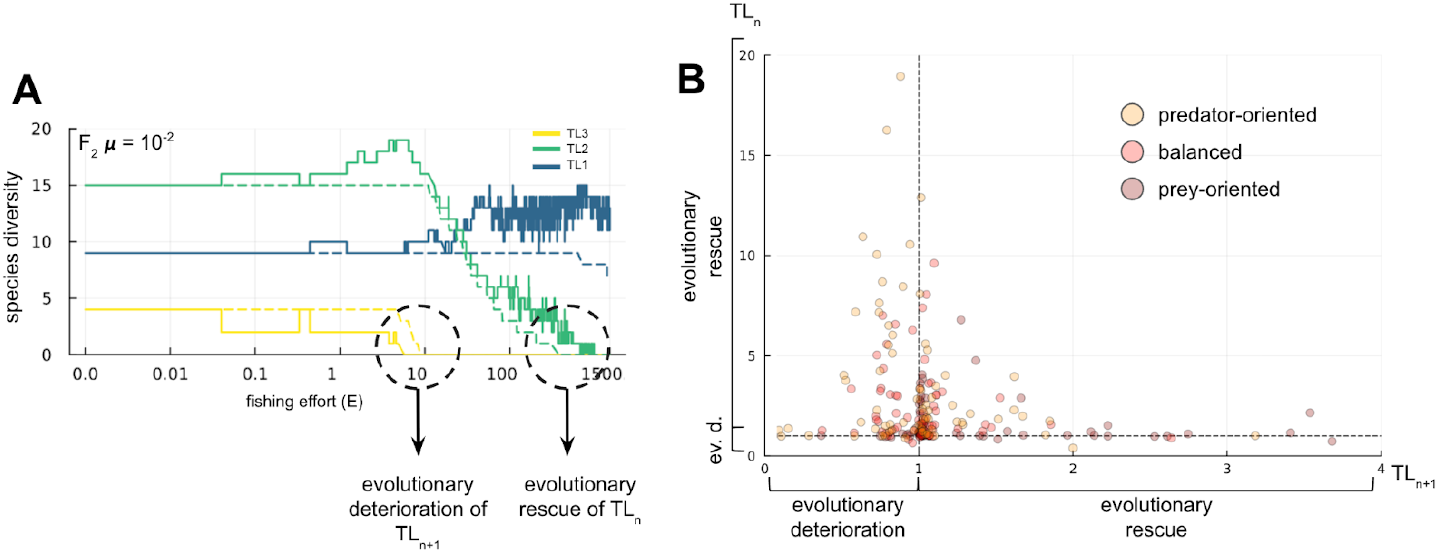
Relationship between evolutionary events at trophic level TLn and at the next higher level TLn+1. Food webs analyzed are *Nw*_1_ with μ = 10^−1^. A: Example on the network shown in Figure 2. B: Intensity of evolutionary rescue (*E*_*v*_ > *E*_0_) or evolutionary deterioration (*E*_*v*_ < *E*_0_), expressed as the ratio *E*_*v*_/*E*_0_, at TLn relative to TLn+1.

